# Predicting the evolution of sexual dimorphism in gene expression

**DOI:** 10.1101/2020.05.26.116681

**Authors:** David Houle, Changde Cheng

## Abstract

Sexual dimorphism in gene expression is likely to be the underlying source of dimorphism in a variety of traits. Many analyses implicitly make the assumption that dimorphism only evolves when selection favors different phenotypes in the two sexes, although theory makes clear that it can also evolve as an indirect response to other kinds of selection. Furthermore, previous analyses consider the evolution of a single transcript or trait at a time, ignoring the genetic covariance with other transcripts and traits. We first show which aspects of the genetic-variance covariance matrix, **G**, affect dimorphism when these assumptions about selection are relaxed. We then reanalyze gene expression data from *Drosophila melanogaster* with these predictions in mind. Dimorphism of gene expression for individual transcripts shows the signature of both direct selection for dimorphism and indirect responses to selection. To account for the effect of measurement error on evolutionary predictions, we estimated a **G** matrix for eight linear combinations of expression traits. Sex-specific genetic variances in female- and male-biased transcription, as well as one relatively unbiased combination were quite unequal, ensuring that most forms of selection on these traits will have large effects on dimorphism. Predictions of response to selection based on the whole **G** matrix showed that sexually concordant and antagonistic selection are equally capable of changing sexual dimorphism. In addition, the indirect responses of dimorphism due to cross-trait covariances were quite substantial. The assumption that sexual dimorphism in transcription is an adaptation is likely to be incorrect in many specific cases.

## Introduction

The prevailing model of the evolution of sexual dimorphism (e.g., Rice and Chippindale 2001; Bonduriansky and Chenoweth 2009; Cox and Calsbeek 2009) supposes that a sexually monomorphic ancestral population is subjected to novel selective pressures that drives the male and female means apart. Examples of extreme sexual dimorphism tied sex-specific functions, such as the horns of bighorn sheep, constitute compelling evidence for this scenario.

Another kind of evidence for this model is the existence of sexual conflict, that is, persistent antagonistic selection on sex-specific traits (Rice 1984; Partridge and Hurst 1998; Arnqvist and Rowe 2005). Since the genomes of the two sexes are extremely similar, differing principally in the sex chromosomes, a population may evolve towards the sex-specific optima very slowly (Lande 1980), resulting in sexual dimorphism that is less than optimal. There is direct experimental evidence in favor of such conflicts from experiments that alter the relative strength of selection on the sexes, and result in changes in traits expressed in both sexes (Prasad, et al. 2007). We call this the genomic constraint hypothesis of sexual dimorphism.

A readily available source of high-throughput data on sexual dimorphism is messenger RNA abundance in the sexes (Ingleby, et al. 2015; Mank 2017). Furthermore, differences in gene expression are likely to underlie dimorphism in many other phenotypic traits. Several lines of evidence suggest that genomic constraint and intra-locus conflict shapes the evolution of gene expression in *Drosophila melanogaster*. Hollis, et al. (2014) minimized sexual conflict by enforcing monogamous matings for more than 100 generations, and observed that expression of sex-biased transcripts shifted in the direction of female expression. This suggests that biased genes are on average less different in their expression than would be optimal. Griffin et al. (2013) reanalyzed the gene expression data of Ayroles et al. (2009), and found that estimates of genetic correlations between male and female gene expression for a particular gene, *r_MF_*, were correlated with multiple aspects of sexual dimorphism, including the degree of sex bias within *D. melanogaster*, the rate of evolution of expression bias among species, and the degree of sexually antagonistic selection that *D. melanogaster* experiences. In addition, a previous study of gene expression identified transcripts with sex by fitness interactions that potentially indicate antagonistic selection (Innocenti and Morrow 2010), and these transcripts exhibited larger *r_MF_*.than other transcripts. All of these results are expected under the genomic constraint hypothesis.

On the other hand, concordant selection on the sexes can also result in increases in dimorphism if the sexes differ in their evolvability (Fisher 1930; Lande 1980; Leutenegger and Cheverud 1982; Cheverud, et al. 1985; Lynch and Walsh 1998, Chapter 24; Bonduriansky and Chenoweth 2009; Wyman, et al. 2013). The model of Connallon and Clark (2014) elegantly combines the effects of antagonistic and concordant selection. It shows that under general conditions almost any change in the sex-specific optima will generate at least transient dimorphism and sexual conflict even when selection on the two sexes is initially concordant. Despite the widespread acknowledgement by theoreticians of concordant selection’s possible role in the evolution of dimorphism, analyses of empirical data rarely incorporate this possibility. The analysis of gene expression by Griffin et al. (2013) considered did not consider any alternatives to the genomic constraint hypothesis.

A quantitative genetic framework is useful to capture the nature of the genetic variation currently segregating within populations to either allow or constrain alterations in the level of sexual dimorphism (Lande 1980). In this framework the variances and covariances among male- and female-expressed traits, summarized in a **G** matrix, make it possible to predict how traits will respond to current selection. The covariances between trait values in one sex with those in the other sex are key to the potential resolution of sexual conflicts. These covariances can be collected into a sub-matrix of **G** known as the **B** matrix. The diagonals of the **B** matrix are the genetic covariances of homologous traits expressed in different sexes. They are commonly summarized using the genetic correlation *r_MF_*.

We set out to expand on the analyses of Griffin et al. (2013) because they made simplifying assumptions about the genetic context in which dimorphism evolves, and their analysis does not fully match the nature of the data. From a genetic perspective, there is potentially more to the evolution of sexual dimorphism than is captured by *r_MF_*. As noted above, differences in genetic variances between the sexes can also generate dimorphism. In addition, gene expression of each transcript genetically covaries with a vast array of other transcripts, as well as other kinds of phenotypes. Indeed, Ayroles et al. (2009), the workers who collected the data reanalyzed by Griffin et al., detected 241 clusters of transcripts that were positively correlated with transcripts in that cluster, and more independent of, or even negatively correlated with expression of transcripts in other clusters. These covariances may cause a focal trait and its dimorphism to evolve as an indirect response to selection (Blows and Hoffmann 2005; Hansen and Houle 2008; Walsh and Blows 2009).

Turning to the statistical aspects of the data, there are two key issues with the Griffin et al. (2013) analyses. First, Griffin et al. treated each estimate of *r_MF_* as statistically independent, despite the evidence of among-transcript correlations. Second, estimates of *r_MF_* are biased downwards due to sampling variance, as noted by Griffin et al. (2013). They demonstrated the direction and degree of bias by analyzing subsets of transcripts with different repeatabilities, but did not make any unbiased estimates of *r_MF_*.

We reanalyzed the Ayroles et al. (2009) expression data set using two approaches. We first analyzed the entire sample covariance matrix of line means, which we term **G**\*. **G**\* allows us to investigate whether the cross-transcript and cross-sex covariances can explain more variation in sexual dimorphism than 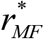 alone. This is the same approach as that of Griffin et al. (2013). To overcome the lack of independence in the estimates, we made inferences based the distributions of bootstrapped estimates. Second, we estimated a **G** matrix for a small number of linear combinations of gene expression traits that collectively have statistically significant genetic variances. This yields a statistically unbiased **G** matrix, including estimates of *r_MF_*, which we use to compare the predicted selection responses to antagonistic and concordant selection. Before proceeding to these results, we first build intuition about the roles of asymmetries in male and female genetic variance, and of cross-trait covariances in the evolution of sexual dimorphism.

### What influences the rate of evolution of sexual dimorphism?

Consider *k* quantitative traits, with phenotypic values *z_1_, z_2_,…, z_k_*. Lande (1979) formulated the quantitative genetic prediction equation

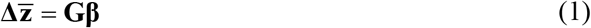

that predicts a *k* x 1 vector of predicted responses to selection, 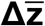, from a *k* x 1 vector of partial regression coefficients of fitness on trait, **β**, and the *k* x *k* additive genetic covariance matrix, **G**. Lande (1980) then generalized this to consider the same *k* traits separately in each sex

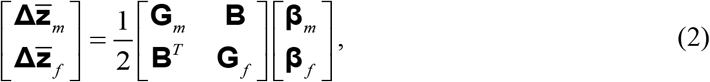

where *f* and *m* index female and male response vectors, **G** matrices, and selection gradient vectors. Individual elements of the **G** matrices will be written *m_ij_* and *f_ij_*. The *k* x *k* matrix **B** contains the covariances between traits expressed in the other sex, with *b_ij_* denoting the covariance of the *i*th trait in females with the *J*th trait in males. The diagonal of **B** is the covariance of homologous traits between the sexes, while the off-diagonal elements are the covariances between sexes among non-homologous traits. The **B** matrix is not necessarily symmetric, as the alleles may have different magnitudes of effects on each sex.

### One trait case

Griffin et al.’s (2013) analysis focused on the genetic correlation between male and female expression for the homologous, *r_MF_*, as an indicator of the degree of constraint. To see the biological meaning of *r_MF_*, consider the simple case of symmetrical antagonistic selection on the expression of one transcript. Under symmetrical antagonistic selection the sign of the selection gradients in the two sexes are reversed. Using the prediction of the response to selection based on **G** we obtain

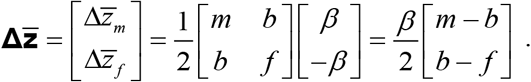

The change in sexual dimorphism is

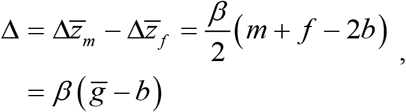

where 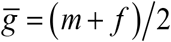 is the average genetic variance in the two sexes. This change can either increase or decrease sexual dimorphism if selection is on a previously dimorphic trait.

The effect of *r_MF_* can be highlighted by substituting 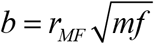, which yields

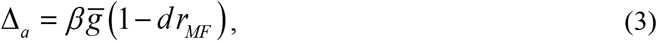

where

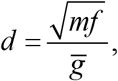

the ratio of the geometric and arithmetic means of the male and female variances. This ratio reaches a maximum value of *d*=1 when the ratio of male to female variances is 1. A positive *r_MF_* restricts the rate of change in sexual dimorphism, while a negative correlation increases it. The sign of the change in dimorphism is controlled by the sign of *β*, as both of the other terms must be positive. Due to the similarity of male and female genomes, we expect that *r_MF_* will generally be positive, restricting the evolution of dimorphism. Reviews of empirical estimates of *r_MF_* confirm that they are positive on average (Poissant, et al. 2010; Griffin, et al. 2013). In the special case when *m*=*f* we recover the familiar result that

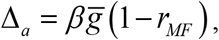

the starting point for Griffin et al.’s analysis. This is, however, also the state where the influence of *r_MF_* is maximized, as *d*<1 whenever *m* ≠ *f*. This effect is relatively small when the ratio of *m* to *f* is fairly close to 1; for example, when the ratio is 2:1 or 1:2, *d* is 0.94; *d* is not halved until the ratio of variances is nearly 14:1.

Under concordant selection, the change in dimorphism is

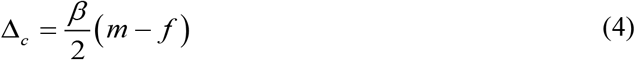

This can be put in the same form as (3), yielding

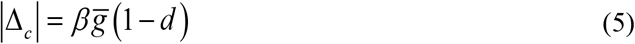

Thus concordant selection will change dimorphism whenever there are different genetic variances in the two sexes, as previously noted by many authors (see the Introduction). Comparison of equations (3) and (5) shows that in the single trait case, the rate of change in dimorphism will be higher under antagonistic selection than concordant selection of equal strength, regardless of the asymmetry in sexspecific variances.

### Two trait case

To get a sense for how other traits could influence these results, we consider a second trait, so *k*=2. In this case, differences in genetic variances between the sexes remain a key source of dimorphism. However, to focus on other features of the **G** matrix that can promote dimorphism, but are absent from the *k*=1 case, we consider the special case when all trait variances are 1, yielding the **G** matrix

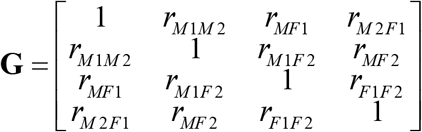

If only the focal trait is under selection (e.g., for antagonistic selection 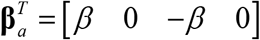, the above results hold for the selected trait. However, there is also an indirect response, which leads to a change in dimorphism in trait 2 under antagonistic selection of

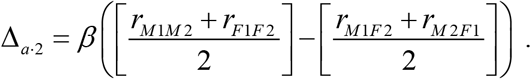

The first term in brackets is the average correlation between the traits in males and females, and the second is the average cross-sex correlation. Under concordant selection, sexual dimorphism changes a the rate

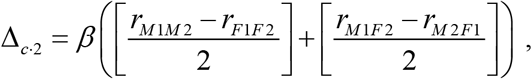

which is affected by the asymmetry of the cross-trait correlations between **G**_m_ and **G**_f_ (the first term in brackets), and between the off-diagonal elements of **B** (the second bracketed term).

The fact that there are indirect responses to selection makes the interpretation of the existing degree of dimorphism in particular traits more challenging. For example, it is quite possible for the change in dimorphism of the selected trait to be less than that of the unselected trait. The direct response to antagonistic selection, Δ_*a*·1_, will be small when *r_MF1_* ≈ 1, making it plausible that Δ_*a*·2_ > Δ_*a*·1_ if within-sex, cross trait correlations *r_M1M2_* and *r_F1F2_* are larger than cross-sex cross-trait correlations *r_M1F2_* and *r_M2F1_*.

If both traits are under directional selection, there are two orthogonal, symmetrically antagonistic selection vectors 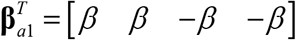 and 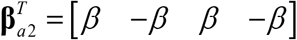. vector **β**_*a*l_, for example, predicts a response vector

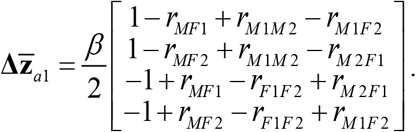

and a change in dimorphism of

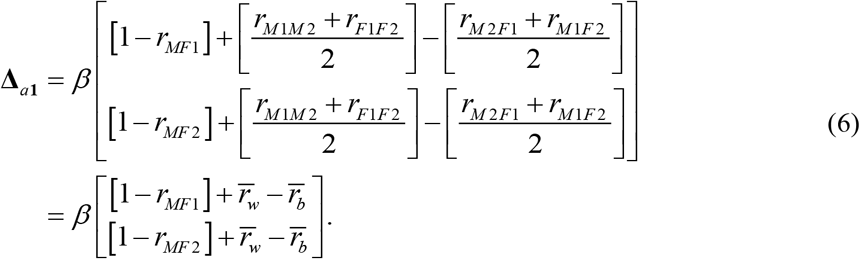

The direct responses are given by the leftmost terms in brackets. The middle terms give the indirect responses due to the average within-sex correlations between traits, 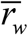. The final term gives the indirect responses due to the average off-diagonal (between-sex) correlations in **B**, 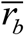.

The complementary antagonistic selection gradient, **β**_*a*2_, reverses the signs of the indirect effects, so that 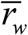 retards the evolution of dimorphism, while 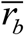 facilitates it. Thus the effects of 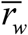 and 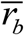 are always antagonistic to each other. Dimorphism is always constrained by *r_MF_*.

In contrast, concordant selection, 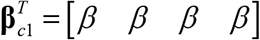, predicts a change in dimorphism of

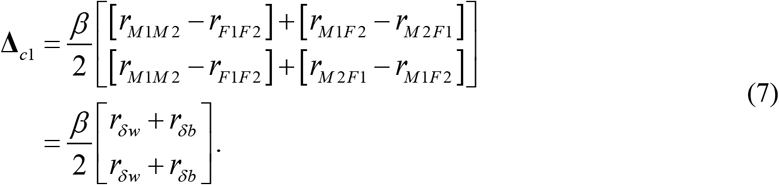

In this special case where all genetic variances are equal there is no direct response in dimorphism. The first term in brackets, *r_δw_*, is the indirect response in asymmetry due to differences between male and female within-sex correlations; the second term, *r_δb_*, is due to asymmetry of the off-diagonal between-sex, between-trait correlations within the **B** matrix. Changing the direction of the concordant vector reverses the sign of the effects of the differences.

Under both antagonistic and concordant selection, the effect of these contrasts vary depending on the nature of selection, and can either enhance or retard the evolution of dimorphism. The overall rate of change in sexual dimorphism is a function of all the components of the **G** matrix. Unlike in the *k=1* case, it is possible for the rate of increase in dimorphism due to concordant selection to exceed that due to antagonistic selection when the strength of selection is held constant.

## Materials and Methods

### Gene expression data

We reanalyzed the adult gene expression data (Ayroles, et al. 2009) gathered for inbred lines of the Drosophila Genome Reference Project (Mackay, et al. 2012). Each DGRP line was independently derived from a different inseminated female *D. melanogaster* sampled from a single outbred population, followed by 20 generations of brother-sister mating.

Ayroles et al. (200) assayed whole body gene expression using Affymetrix Drosophila Genome 2.0 microarrays. They assayed expression twice in each sex in each of 40 DGRP line, for a total of 160 chips. The raw data were downloaded from https://www.ebi.ac.uk/arrayexpress/experiments/E-MEXP-1594. The data were normalized and processed using the R Bioconductor package *oligo* (Carvalho and Irizarry 2010; Huber, et al. 2015). Probe-to-gene mapping was downloaded from https://metazoa.ensembl.org/index.html via BioMart. In some cases, a single probe assayed expression of transcripts at more than one gene. We assigned results from such probes to just one of these genes, chosen arbitrarily. This left expression data on 12,701 genes. Gene expression was in log_2_ units, so differences in expression are equivalent to log_2_(ratios).

### Gene by gene analyses

We tested for the presence of significant genetic variation at each gene using a mixed model analysis with sex, probe, and probe by sex effects fixed, and line, line by sex and line by probe effects treated as random. For 11,039 genes, a single probe was assayed, and the probe effects were omitted from the model. We compared the likelihood of the data for models that included or omitted all the random effect terms using a likelihood ratio test with two df for the genes with only one probe, and three df for genes with more than one probe. Genes were retained for further analysis if *P*<0.01 for the total line effects. Mixed model analyses were fit using Proc Mixed in SAS/STAT software (SAS Institute 2016).

For genes with significant line effects, we retained the least squares means from this model for each sex and line, and then calculated the covariance matrix of these means to form **G**\*. Sexual dimorphism in expression of the *i*th gene, 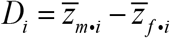, was measured as the difference between the least squares means of male and female log_2_-transformed gene expression. We defined three categories of sex bias: male-biased (MB) genes with *D_i_* > 1, female-biased (FB) genes with *D_i_ <* −1, and relatively unbiased (UB) genes with |*D*| ≤ 1.

From equation (3) we predict that if antagonistic selection plays a major role, |*D_i_*| will be positively related to 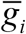, and negatively related to *r_MF·i_*. From equation (4) we predict a positive relationship between *|m_i_ – f_i_|* and |*D*| under concordant selection. From equation (6), we can see that the averages of the corresponding off-diagonal elements of 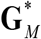 and 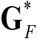, and of off-diagonal elements above and below the diagonal of **B\*** affect the evolution of dimorphism under antagonistic selection. To capture the effect of within-sex correlations for the *i*th gene, we calculated

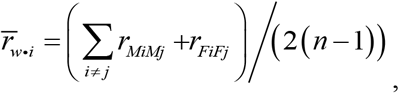

where the summation is over all genes, and *n* is the number of genes. To summarize the effects of between-sex correlations we used

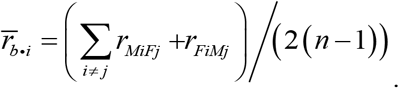

From equation (7), we can see that differences of the corresponding off-diagonal elements of 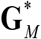 and 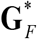, and of off-diagonal elements above and below the diagonal of **B\*** promote the evolution of dimorphism under concordant selection. We quantified the effects of the differences in within-sex correlations as

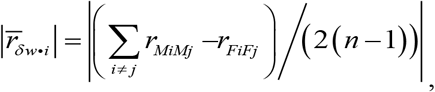

and the differences in between-sex correlations as

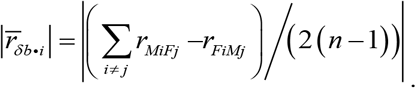

In addition, we also investigated the effects of the mean level of expression, 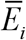, and an index of the tissue specificity of gene expression, *τ_i_* (Yanai, et al. 2004), on the level of sexual dimorphism. The tissue specificity index ranges from 0 when all tissues have equal expression, to 1 when one tissue accounts for all expression. We downloaded tissue-specific expression data from http://flyatlas.org/ Geo accession GSE7763, and calculated *τ* for each transcript. The value of *τ_i_* is the average *τ* over all transcripts of gene *i*.

Values of |*D*| must be positive, but are strongly right skewed with a small minority of very large values. To reduce the influence of this large tail, while preventing values very near 0 from causing a left-skew to the transformed data we transformed dimorphism as log_10_ (|*D_i_*| + 0.01). We log 10 transformed 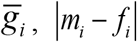 and 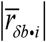 before analysis for similar reasons.

### Selection on gene expression in *Drosophila melanogaster*

We used results from Innocenti and Morrow (2010) to investigate whether evidence of laboratory selection on transcript abundance helps explain dimorphism. Innocenti and Morrow studied sex-specific fitness of 100 heterozygous haploid genotypes in males and females, and found a significant negative genetic correlation between fitness in male and female genotypes. They then estimated adult gene expression for 15 of these genotypes that spanned the range of male and female fitnesses using Affymetrix Drosophila Genome 2.0 arrays. They tested whether fitness predicted the expression of each transcript, and for a sex by fitness interaction. They report *P*-values for tests judged to be significant in their Table S1, so it is unclear how many total transcripts were tested. A total of 608 transcripts had a significant main effect, 1478 had a significant interaction, while 392 had both a main and a significant interaction effect. We were able to associate these significant transcripts with a larger number of genes than Innocenti and Morrow did, presumably because of increased annotations since that analysis was undertaken. When paired with the Ayroles et al. (2009) data set, 516 genes had least one transcript with a significant main effect, 1300 with a significant interaction, and 348 for which both effects were significant.

### Quantitative genetic analysis of bias classes

To generate a small number of variables that summarized the variation in expression, we performed separate principal component analyses (PCA) on the covariances of least-squares means of expression in each sex for the male-, female, and relatively-biased genes. For the two biased sets, we performed a PCA of just the expression of the dominant sex. For the unbiased genes, we performed a PCA on the covariance matrix of line-sex averages. Multiplying the sex-specific mean data by the eigenvectors computes a weighted average of the expression of the genes in each bias subset, resulting in scores that are defined in the same manner in each sex.

We partitioned the variance in the principal component scores into genetic and non-genetic sources using restricted maximum-likelihood implemented in the program Wombat (Meyer 2006-2019). We assumed that all the inbred lines were unrelated. Whole-genome sequencing of these lines suggests that this is a good, albeit imperfect, approximation of the relationship among them (Huang, et al. 2014). Estimation of the genetic variance-covariance matrix, **G**, was carried out for both full- and reduced-rank models (Kirkpatrick and Meyer 2004; Meyer and Kirkpatrick 2005, 2008), and we selected the bestfitting model on the basis of Akaike’s information criterion corrected for small sample size (AICc).

We assessed the fit of models with different numbers of traits drawn from the three expression bias classes, and eventually attempted full analyses of 16 traits, two expression vectors from each of the female-biased and male-biased gene sets, and four from the relatively unbiased class. We were unable to estimate the average information matrix for this 16-trait matrix. We traced this to a lack of significant genetic variation in the female expression of male-biased genes or in the male expression of female-biased genes, which were then dropped from the analysis, resulting in a 12-trait data set. Sampling variances of matrix elements remained stable under models fit to even fewer traits. Once we obtained well-estimated matrices, Results we back-transformed estimates to the original scores and used the REML-MVN approach (Meyer and Houle 2013; Houle and Meyer 2015) to generate 1,000 replicate matrices drawn from the sampling distribution of the matrix **G**. These replicate estimates of **G** were used to generate the sampling distribution of the predicted responses to selection. In most cases, the distribution of **G**-derived statistics were asymmetrical, so we report medians and 2.5% and 97.5% quantiles.

### Analyses of modified G matrices

To investigate which aspects of **G** matrix structure have effects on the evolution of dimorphism, we formed five modified matrices that highlight those aspects of **G** that affect the evolution of dimorphism. The *k*=2 theory developed above, as well as the more general analysis of Cheng and Houle (submitted) suggests that the evolvability of dimorphism to antagonistic selection is promoted the average genetic variance in males and females, 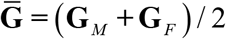, and restricted by the symmetrical component of **B**, **B**_*5*_ = (**B + B**^*T*^)/2. When these matrices are substituted for **G**_*M*_, **G**_*F*_, **B** and **B**^*T*^ this eliminates the components of **G** that cause evolvability of dimorphism under concordant selection, namely the asymmetry between **G**_*M*_ and **G**_*F*_, (**G**_*M*_ - **G**_*F*_) / 2 and the asymmetrical component of **B**, **B**_*A*_ =(**B** - **B**^*T*^)/2. In addition we explored the effect of eliminating **B** by replacing it with the zero matrix **0**.

## Results

### Gene-wise analysis

Of the 12,071 genes investigated, we detected significant genetic variation at P<0.01 in 10,489 genes, similar to the results obtained by Ayroles et al. (2009). We do not consider the non-significant genes further. Of the significant genes, 1,447 were male biased (MB: *D_i_ >* 1, a ratio of male to female expression greater than 2), 2,073 were female biased (FB: *D_i_* < −1, a ratio less than 0.5), and the remaining 6,979 genes had relatively unbiased expression (UB: |*D*| ≤ 1, within a factor of two between the sexes).

### Which aspects of G* correlate with dimorphism of transcription?

Under the familiar hypothesis that sexually antagonistic selection (SAS) drives the evolution of dimorphism, equations (3) and (6) predict that the absolute value of sexual dimorphism, |*D*|, will be positively related to 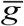 and negatively related to *r_MF_*. In addition, 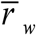 and 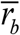 have conflicting effects on the change in dimorphism. Their net effect can be positive or negative. If sexually concordant selection (SCS) influences dimorphism, equation (4) shows that differences in male and female genetic variances should be positively correlated with |*D*|, while equation (7) predicts that the absolute value of the average differences of within- 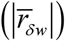 and between-sex correlations 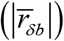 should also be positively correlated with *|D|*. In addition previous analyses suggest that the mean level of gene expression, 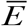 and the degree of tissue-specifity, *τ*, are also related to |*D*|.

Figure 1 shows the relationships of log_10_ (|*D*| + 0.01) for all 10,489 genes with these **G**-matrix predictor variables. The Pearson correlation matrix of these plus the average level of expression, 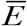, and an index of tissue specificity, *τ*, is shown in Table S1. The predicted relationships of *r_MF_* and log_10_ 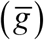 under SAS are clearly evident. In addition, 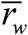 has a strong positive relationship with log_10_ (|*D*| + 0.01), suggesting the effects of the within-sex between-trait correlations swamp those of the between-sex between-trait correlations captured by 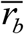. It is evident from Figure 1 that the average value of 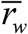 is larger than that of 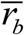. Both log_10_ (*g_m_ - g_f_*) and 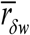 have clear positive relationships with dimorphism as predicted under SCS.

**Figure 1.**
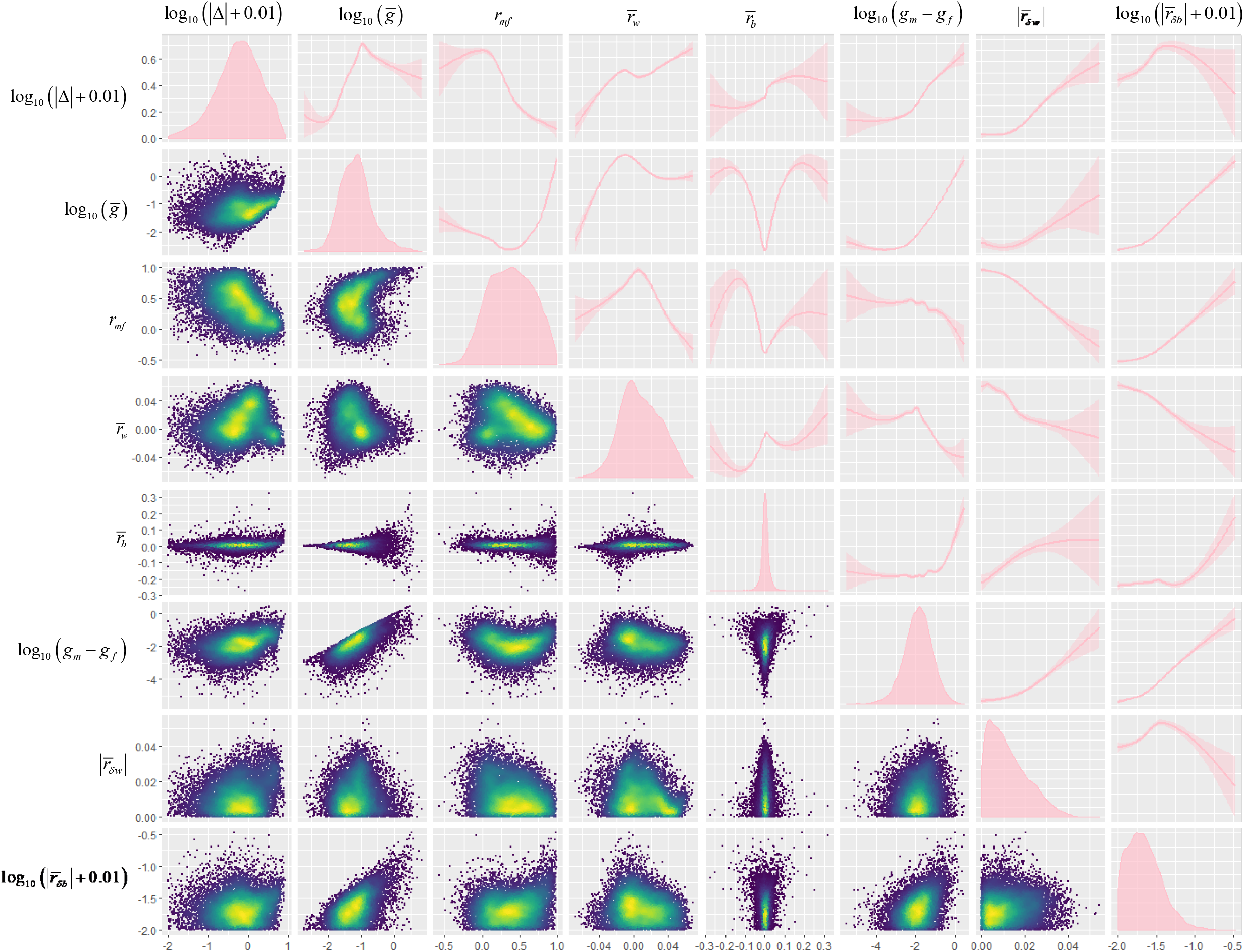
Distribution, density and smoothed trend plots for log_10_ (|Δ| + 0.01) and potentially predictive aspects of the **G** matrix. Visualization prepared in the ggplot2 package in R (R Core Team 2020), using default smoothing and density plot parameters

These raw relationships are confounded, as Figure 1 and Supplementary Table S1 also show that these predictor variables are all correlated with each other. Log_10_ (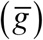) and log_10_ (*g_m_ - g_f_*) are particularly strongly correlated. In addition, some of the relationships are strongly non-linear. Table S1 shows that *τ* is positively correlated with the measures of genetic variation. These features make interpretation of the raw relationships difficult. Furthermore, the expression of individual genes are not independent. To disentangle these factors, we used multiple regression of log_10_ (|*D*| + 0.01) on the other variables, bootstrapped at the level of inbred lines. The results are shown in Table 1. When all genes are included, the model explains 40% of the variation in dimorphism. Both mean expression 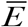 and tissue specificity *τ* have consistent positive effects on dimorphism, as shown by the positive signs of the bootstrap quantiles. Log_10_ 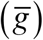, *r_mf_* and 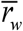 have relationships with log_10_ (|*D*| + 0.01), explaining 2.1, 15.5 and 6.7% of the variance respectively. In addition, there is also a consistent positive relationship of log_10_ (| *m* — *f* |) with log_10_ (|*D*| + 0.01) as expected under SCS, explaining 1.1% of the variance. The effects of the differences of within-sex and between-sex correlations have no significant effects. Expression properties 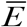 and *τ* also have significant positive relationships with log_10_ (|*D*| + 0.01), explaining 3.2 and 3.5% of the variation.

**Table 1.** Regression of log_10_ (|*D*| + 0.01) on expression characteristics.

| Parameter<sup>a</sup> | Pred.<sup>b</sup> | All genes |  |  | Biased<sup>d</sup> |  |  | Unbiased<sup>d</sup> |  |  |
| --- | --- | --- | --- | --- | --- | --- | --- | --- | --- | --- |
|  |  | Median | Quantiles<sup>c</sup><br>2.5%, 97.5% | R<sup>2</sup> | Median | Quantiles<sup>c</sup><br>2.5%, 97.5% | R<sup>2</sup> | Median | Quantiles<sup>c</sup><br>2.5%, 97.5% | R<sup>2</sup> |
| Mean expression $\bar{E}$ | | **0.04** | **0.03, 0.05** | **3.2** | **0.03** | **0.02, 0.03** | **6.2** | **0.03** | **0.03, 0.04** | **1.8** |
| $\tau$ | | **0.78** | **0.49, 0.96** | **3.5** | **0.99** | **0.82, 1.12** | **32.6** | -0.01 | -0.10, 0.10 | 0.0 |
| $\log_{10}(\bar{g})$ | A+ | **0.20** | **0.02, 0.34** | **2.1** | 0.05 | -0.04, 0.11 | 0.8 | **0.14** | **0.10, 0.19** | **1.2** |
| $r_{mf}$ | A- | **-0.63** | **-0.76, -0.48** | **15.5** | **-0.10** | **-0.15, -0.04** | **4.5** | **-0.28** | **-0.38, -0.19** | **2.3** |
| $\bar{r}_w$ | A? | **4.81** | **2.99, 7.66** | **6.7** | **0.41** | **0.19, 0.60** | **0.3** | -0.95 | -2.13, 0.50 | 4.2 |
| $\bar{r}_b$ | A? | 0.03 | -1.52, 1.41 | 0.1 | 0.09 | -0.08, 0.30 | 0.2 | 0.15 | -0.49, 0.96 | 0.0 |
| $\log_{10}(\vert m - f\vert )$ | C+ | **0.11** | **0.07, 0.16** | 1.1 | 0.00 | -0.02, 0.02 | 0.3 | **0.04** | **0.02, 0.07** | **0.4** |
| $\vert \bar{r}_{\delta w}\vert $ | C+ | 3.72 | -6.10, 14.23 | 1.8 | 0.01 | -0.44, 0.52 | 1.3 | 1.17 | -1.58, 5.59 | 0.6 |
| $\log_{10}(\vert \bar{r}_{\delta b}\vert + 0.01)$ | C+ | -0.09 | -0.23, 0.20 | 0.0 | -0.00 | -0.08, 0.13 | 0.4 | -0.07 | -0.14, 0.01 | 0.1 |

<sup>a</sup>Parameter symbols explained in the text; $\bar{r}_w$ and $\bar{r}_b$ generalize the indirect selection parameters defined in equation (6) to include the mean of the average within-sex correlations of all traits with expression of the focal gene, while $|\bar{r}_{\delta w}|$ and $|\bar{r}_{\delta b}|$ generalize those in equation (7) to include the absolute values of the mean differences of all within-sex correlations with the focal gene, or the differences in the between-sex correlations involving the focal gene. $|\bar{r}_{\delta b}|$ was log-transformed to minimize the influence of observations with exceptionally high values.
<sup>b</sup>Predicted sign of relationships with $\log_{10}(|D| + 0.01)$ under concordant or antagonistic selection; A=antagonistic prediction,
C=concordant prediction; ? indicates that both $\bar{r}_w$ and $\bar{r}_b$ are predicted to affect dimorphism under SAS, but the sign of these effects depends on the details of selection.
<sup>c</sup>Quantiles from 1000 bootstrap resamples at the inbred line level. When the bootstrap 95% quantiles have consistent sign, we consider the effects to be statistically significant. Significant values are shown in bold.
<sup>d</sup>Biased genes have $|\Delta| > 1$ ; Unbiased genes $|\Delta| \leq 1$ .

Separate analyses of the biased (|*D*| > 1) genes shows that the **G** matrix properties are much less predictive of dimorphism in this subset. 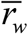 and *τ* remain significant, and the role of *τ* is much stronger in this class of genes, explaining 32.6% of the variation. Of the **G** matrix properties, *r_mf_* and 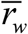 remain consistent predictors of dimorphism, although the variance explained is much reduced from that over the entire data set. Analysis of just the relatively unbiased (|*D*| ≤ 1) genes shows that 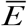 is again a significant predictor, but *τ* is not. **G** matrix properties expected to covary with log_10_ (|*D*| + 0.01) under both SAS and SCS are significant in this subset; the relationships with log_10_ 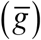 and *r_mf_* suggest the action of SAS, while the positive relationship with log_10_ (|*m* - *f*|) is consistent with SCS. All of these variables explained a small proportion of the variance in dimorphism. On balance, these results indicate a strong signature of SAS over the entire range of dimorphism values, and effects of SCS on dimorphism that is less prevalent on genes with high levels of dimorphism.

Innocenti and Morrow (2010) estimated selection on expression for a subset of the transcripts in our data set. They treated significant sex by fitness interactions for transcription (their models used transcription as the dependent variable) as equivalent to SAS. We use the fact that the partially overlapping set of transcripts with a significant main effect of fitness experience SCS. In general, any selection will include both concordant and antagonistic components (Cheng and Houle accepted). Unfortunately Innocenti and Morrow (2010) did not report selection effect sizes, so their results can only tell us whether there was at least some evidence of SCS and SAS for each transcript. When we entered indicator variables for at least one significant transcript main-effect and sex-by-fitness effects into the multiple regression model, both indicators had a small but significant negative effect on |*D*| (results not shown). Perhaps more importantly, the magnitude of these effects was small, and the parameter estimates shown in Table 1 changed little when the indicators of selection were included in the multiple regression model. There is no clear predicted relationship between ongoing selection and dimorphism. One potential explanation for the negative relationship between selection and dimorphism is that Innocenti and Morrow’s study had greater power to detect selection in less dimorphic transcripts.

### Quantitative genetic analysis of gene expression

Ayroles et al. (2009) measured gene expression in just 40 inbred lines, which precludes estimating a G matrix for more than a small number of traits. To choose informative traits, we first calculated raw among line covariance matrices, G*, for each of the three bias classes: male expression of male biased genes (MB), female expression of female biased genes (FB), and the sex-averaged expression of the remaining genes, termed relatively unbiased (UB). We then conducted a principal components analysis of each G*. Male and female expression of each line was then scored on the first two PCs in the MB and FB subsets, and the first four in the UB subset, so *k* = 8 in each sex.

Analyses of genetic variation of the *k* = 8 data showed that the best fitting model had eight variable dimensions vectors out of 16 possible (and fit more than 3.6 AICc units better than the 7- or 9dimensional models). The covariance and correlation matrices estimated using this model are shown in Supplementary Table S2. The average information matrix contained very small elements that prevented estimation of standard errors for all model terms, suggesting that the overall fit of the model was poor.

To help diagnose the trait combinations lacking significant genetic variation, we examined the estimates of trait variances, shown in Supplementary Table S3. Sex-specific variances are substantially different for the biased expression classes (MB and FB). Note that these biased traits as well as trait UB1 have very large asymmetries in the sex-specific genetic variances, making *d* (equation 3) substantially less than 1. This indicates that dimorphism is substantially more likely to evolve under both SAS (equation 3) and SCS (equation 4) than under the assumption of equal variances.

We conjectured that the female variances for the male-based traits and the male variances for the female-biased traits are probably not distinguishable from 0, judging by the fact that these estimates are less than 2.5 standard errors (S.E.) from 0 (Sztepanacz and Blows 2017). To test this, we dropped the female expression of male-biased traits and the male expression of female-biased traits from the data set, and repeated estimation of the genetic variance for the remaining 12 variables. The best model now had nine significant genetic dimensions, although this was just 0.91 AICc units better than the eight-dimensional model. This supports the hypothesis that there is no significant genetic variation in females for male-biased genes or in males for female biased genes. We predict that direct selection to alter expression in the low-expressing sex will be relatively ineffective.

The resulting genetic correlation and covariance matrices from the twelve-trait, nine-dimensional model are shown in Table 2. Note that while we have retained six traits in each sex, only the relatively unbiased expression traits (UB1-UB4) are shared between the sexes. Thus, only the portions of Table 2 involving these traits (outlined in boxes) correspond to Lande’s (1980) partition of the **G** matrix. The intersexual correlations for these homologous traits, *r_MF_*, shown in bold in the upper right-hand box, average 0.82. This is a little higher than the average value of 0.75 found in a review of previous studies (Poissant, et al. 2010), and much higher than the average *r_MF_* value of estimated in Griffin et al. (2013). All *r_MF_* are greater than 0.5, and traits UB2-4 have *r_MF_* > 0.85. All three of these correlations are less than 3 S.E. from the perfect correlation of 1, suggesting that there may be very little variation to generate conflicting responses in the sexes for these traits (Sztepanacz and Blows 2017).

**Table 2.** Genetic correlation and covariance matrices from the 12 trait analysis. Genetic variances are shown in bold on the main diagonal. Genetic correlations are above the main diagonal, and genetic covariances are below. Sampling standard errors are shown in italic font on the line below the estimates. Boxes outline the submatrices that form the **G** and correlation matrices for the four relatively unbiased traits. The upper right and lower left boxes outline the - matrices of correlations and covariances between homologous trait in males and females, with the diagonal elements shown in bold italics.

|  |  | Male Expression |  |  |  |  |  | Female Expression |  |  |  |  |  |
| --- | --- | --- | --- | --- | --- | --- | --- | --- | --- | --- | --- | --- | --- |
|  |  | MB1 | MB2 | UB1 | UB2 | UB3 | UB4 | FB1 | FB2 | UB1 | UB2 | UB3 | UB4 |
| Male | MB1 | **55.3** | -0.17 | -0.45 | -0.53 | -0.36 | -0.02 | -0.12 | -0.01 | 0.10 | -0.50 | -0.15 | -0.02 |
|  |  | *15.1* | *0.19* | *0.15* | *0.13* | *0.16* | *0.18* | *0.19* | *0.18* | *0.18* | *0.15* | *0.18* | *0.18* |
| Expression | MB2 | -5.1 | **16.7** | -0.18 | -0.35 | 0.48 | 0.27 | -0.05 | -0.31 | -0.02 | -0.45 | 0.50 | 0.08 |
|  |  | *5.7* | *4.3* | *0.17* | *0.15* | *0.14* | *0.17* | *0.18* | *0.16* | *0.17* | *0.15* | *0.14* | *0.18* |
|  | UB1 | -15.5 | -3.4 | **21.4** | 0.01 | -0.15 | -0.15 | -0.50 | -0.14 | **0.60** | 0.16 | -0.14 | -0.32 |
|  |  | *6.5* | *3.3* | *5.0* | *0.16* | *0.16* | *0.17* | *0.13* | *0.16* | *0.11* | *0.16* | *0.17* | *0.15* |
|  | UB2 | -29.1 | -10.5 | 0.3 | **53.7** | 0.05 | -0.10 | 0.28 | 0.22 | -0.37 | **0.87** | -0.14 | 0.08 |
|  |  | *10.9* | *5.5* | *5.5* | *12.4* | *0.16* | *0.17* | *0.16* | *0.16* | *0.14* | *0.05* | *0.16* | *0.17* |
|  | UB3 | -17.3 | 12.8 | -4.6 | 2.3 | **42.2** | 0.09 | -0.01 | -0.29 | -0.14 | -0.00 | **0.93** | -0.03 |
|  |  | *9.2* | *5.0* | *5.0* | *7.9* | *10.0* | *0.17* | *0.18* | *0.16* | *0.16* | *0.17* | *0.03* | *0.17* |
|  | UB4 | -0.9 | 6.5 | -4.1 | -4.1 | 3.4 | **34.5** | 0.08 | -0.05 | -0.02 | -0.13 | 0.08 | **0.85** |
|  |  | *7.8* | *4.4* | *4.6* | *7.2* | *6.5* | *8.4* | *0.18* | *0.17* | *0.17* | *0.17* | *0.17* | *0.06* |
| Female | FB1 | -7.9 | -1.9 | -21.4 | 18.8 | -0.6 | 4.4 | **85.3** | 0.02 | -0.96 | -0.07 | -0.26 | 0.02 |
|  |  | *13.1* | *7.0* | *8.3* | *12.1* | *10.5* | *9.7* | *22.3* | *0.18* | *0.02* | *0.18* | *0.17* | *0.18* |
| Expression | FB2 | -0.5 | -7.2 | -3.6 | 8.8 | -10.4 | -1.5 | 0.7 | **31.2** | 0.06 | 0.25 | -0.40 | 0.31 |
|  |  | *7.5* | *4.1* | *4.3* | *6.9* | *6.27* | *5.6* | *9.1* | *7.4* | *0.16* | *0.16* | *0.14* | *0.15* |
|  | UB1 | 7.5 | -0.7 | **28.2** | -27.6 | -9.4 | -0.9 | -90.5 | 3.2 | **103.5** | 0.00 | 0.09 | 0.03 |
|  |  | *13.6* | *7.2* | *8.9* | *12.9* | *11.0* | *10.0* | *21.8* | *9.4* | *23.8* | *0.17* | *0.17* | *0.17* |
|  | UB2 | -21.5 | -10.6 | 4.3 | **36.8** | -0.1 | -4.3 | -3.9 | 8.1 | 0.2 | **33.4** | -0.11 | 0.14 |
|  |  | *8.4* | *4.5* | *4.5* | *9.1* | *6.3* | *5.8* | *9.6* | *5.6* | *9.8* | *8.1* | *0.17* | *0.17* |
|  | UB3 | -6.8 | 12.5 | -3.9 | -6.5 | **37.2** | 3.0 | -14.5 | -13.7 | 5.6 | -4.0 | **38.1** | -0.01 |
|  |  | *8.4* | *4.9* | *4.8* | *7.6* | *8.9* | *6.2* | *10.5* | *6.3* | *10.5* | *6.1* | *9.2* | *0.17* |
|  | UB4 | -0.9 | 2.1 | -9.3 | 3.7 | -1.1 | **31.6** | 1.4 | 10.9 | 1.7 | 5.3 | -0.2 | **39.9** |
|  |  | *8.6* | *4.6* | *5.1* | *7.8* | *7.0* | *8.1* | *10.4* | *6.3* | *10.8* | *6.3* | *6.7* | *9.7* |

Examination of Table 2 also reveals several other correlations indicative of constraints on the evolution of biased gene expression. The most striking of these is female expression of female-biased PC1 (FB1) and unbiased PC1 (UB1) where *r_MF_* = −0.963. This large negative correlation will constrain the response to selection to increase or decrease both FB1 and UB1, or to hold one constant while changing the other. In addition, many of the correlations between male-biased expression traits and male expression of the relatively unbiased traits have absolute values in the neighborhood of 0.4-0.5, as does the correlation of FB2 with female expression of UB3. Most of these correlations are negative, which would tend to facilitate increasing one trait while decreasing the other. Taken together, these elements show the potential for complex constraints between unbiased and biased genes, beyond that expected due to *r_MF_*.

### Predicted responses to selection

To explore the effects of the covariance structure on potential responses to selection, we compared the predicted responses of the UB traits to sexually antagonistic (SAS) and concordant selection (SCS) of equal strength. We predicted the responses to symmetrical antagonistic and concordant versions of five different selection gradients: directional selection on each of the four UB traits, and directional selection on all four simultaneously with the results shown in Table 3.

**Table 3.** Comparative responses of dimorphism to antagonistic (A) and concordant (C) selection of equal strength (||β|| = 1). Values are medians (2.5% - 97.5% quantiles).

| Sel.<sup>a</sup> | Antagonistic<sup>b</sup> | | Concordant<sup>c</sup> | | UB $\ \Delta\ $ <sup>d</sup> | | | Vector correlation M vs. F | |
| --- | --- | --- | --- | --- | --- | --- | --- | --- | --- |
| | *e* | *r* | *e* | *r* | A | C | ratio | A $\Delta\bar{z}_m \cdot \Delta\bar{z}_f$ | C $\Delta\bar{z}_m \cdot \Delta\bar{z}_f$ |
| UB1 | 35.6<br>(21.0-52.9) | 60.6<br>(35.6-89.2) | 90.8<br>(53.1-133.7) | 104.7<br>(62.9-155.0) | 27.4<br>(16.3-40.5) | 31.2<br>(16.0-49.4) | 0.87<br>(0.66-1.26) | 0.27<br>(-0.22-0.60) | 0.83<br>(0.65-0.97) |
| UB2 | 7.9<br>(4.1-12.7) | 24.9<br>(14.0-38.2) | 87.7<br>(55.1-132.0) | 94.1<br>(60.0-140.6) | 10.4<br>(5.7-16.1) | 16.1<br>(6.6-29.7) | 0.63<br>(0.34-1.46) | 0.34<br>(-0.09-0.60) | 0.89<br>(0.73-0.98) |
| UB3 | 4.2<br>(1.9-7.5) | 15.2<br>(6.8-24.9) | 84.4<br>(50.6-125.7) | 88.6<br>(54.0-130.1) | 6.3<br>(2.8-10.6) | 7.5<br>(3.0-16.5) | 0.81<br>(0.29-2.24) | 0.33<br>(-0.30-0.73) | 0.97<br>(0.88-1.00) |
| UB4 | 7.0<br>(3.7-11.5) | 16.2<br>(8.9-25.8) | 73.0<br>(45.2-108.8) | 77.4<br>(48.5-114.3) | 6.2<br>(3.2-10.7) | 7.2<br>(2.3-18.7) | 0.86<br>(0.28-2.85) | 0.39<br>(-0.52-0.80) | 0.97<br>(0.82-1.00) |
| UB | 27.6<br>(17.3-40.4) | 48.9<br>(28.4-73.3) | 70.4<br>(44.7-103.5) | 75.9<br>(48.7-110.6) | 23.8<br>(14.7-34.7) | 13.2<br>(3.8-26.2) | 1.80<br>(0.93-5.42) | 0.10<br>(-0.46-0.46) | 0.92<br>(0.80-0.99) |

<sup>a</sup>Selection regime: symbols indicate trait subject to directional selection, UB=all 4 UB traits are simultaneously selected.
<sup>b</sup>Selected male traits have positive gradients, while female traits negative selection gradients; *e*=evolvability, the response in the direction of selection; *r*= respondability, the total response to selection.
<sup>c</sup>All selected traits have positive gradients in both sexes.
<sup>d</sup>Predicted change in length of dimorphism vector. A, C=total change in dimorphism under A or C selection; ratio= $\|\Delta_A\|/\|\Delta_C\|$

We show two different estimates of the amount of evolution under each selective regime. Evolvability, *e*, is the response in the direction of the selection gradient, while respondability, *r*, is the length of the total response to selection (Hansen and Houle 2008). As expected from the generally positive intersex correlations, the responses to SCS selection are overall larger than the responses to SAS.

The key prediction concerns the total change in sexual dimorphism, the length of the vector of changes in dimorphism, ||Δ||. Surprisingly, ||Δ|| is often larger under SCS than under SAS. While the confidence limits of the ratios of ||Δ|| due to SAS and SCS are not significantly different from 1 for any of the selection gradients investigated. This results contrasts with the generally assumed scenario that sexual dimorphism is the result of direct selection for dimorphism. The reasons for this counterintuitive result can be diagnosed with reference to the last two columns in Table 3. Responses of each sex to SAS are actually positively correlated, rather than negatively correlated as the direction of selection would suggest. Similarly, the responses of each sex to SCS are not perfectly correlated, as the direction of selection would suggest.

We also predicted the response of dimorphism of the relatively unbiased traits to selection on the sex-biased traits MB1, MB2, FB1 and FB2 with results shown in Supplementary Table S4. The total changes in dimorphism under these scenarios are of comparable magnitudes to the changes arising from selection on the UB traits. Indirect responses of dimorphism to selection on sex-biased traits can have important effects on other traits.

The selection scenarios in Tables 3 and S4 assume that there is no selection on the traits not under directional selection. Alternatively, we can assume that all traits not under directional selection are subject to such strong stabilizing selection. This is the conditional selection scenario (Hansen, et al. 2003; Hansen and Houle 2008). The magnitudes of the conditional responses in dimorphism are much less than for the corresponding unconditional scenarios in Table 3, but SAS and SCS again have similar effects on overall dimorphism, as shown in Supplementary Table S5.

The results in Table 4 show how these predictions are affected by the symmetry of **G**. To make these comparisons, we substituted the symmetrical version of **G**_*M*_ and **G**_*f*_ (their average, 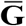) and **B** (**B**_*s*_) into the **G** matrix, and compared the predicted response to those from the unaltered matrix.

**Table 4.** Ratio of changes in relatively unbiased transcript dimorphism (||Δ||) predicted from a modified **G** matrix relative to predictions from the unmodified **G** matrix (results in Table 4).

| $(\text{UB } \ \Delta\ \text{ modified } \mathbf{G}) / (\text{UB } \ \Delta\ \text{ unmodified } \mathbf{G})^b$ | | | | | | | | | | |
| --- | --- | --- | --- | --- | --- | --- | --- | --- | --- | --- |
| Sel.<sup>a</sup> | $\begin{bmatrix} \bar{\mathbf{G}} & \mathbf{B} \\ \mathbf{B}^T & \bar{\mathbf{G}} \end{bmatrix}$ | | $\begin{bmatrix} \mathbf{G}_m & \mathbf{B}_s \\ \mathbf{B}_s & \mathbf{G}_f \end{bmatrix}$ | | $\begin{bmatrix} \mathbf{G}_m & \mathbf{0} \\ \mathbf{0} & \mathbf{G}_f \end{bmatrix}$ | | $\begin{bmatrix} \bar{\mathbf{G}} & \mathbf{B}_s \\ \mathbf{B}_s & \bar{\mathbf{G}} \end{bmatrix}$ | | $\begin{bmatrix} \bar{\mathbf{G}} & \mathbf{0} \\ \mathbf{0} & \bar{\mathbf{G}} \end{bmatrix}$ | |
|  | A | C | A | C | A | C | A | C | A | C |
| UB1 | 1.00 | 0.37 | 1.00 | 0.93 | 1.81 | 0.91 | 1.00 | 0.00 | 1.68 | 0.00 |
| UB2 | 1.00 | 0.77 | 1.00 | 0.53 | 2.53 | 0.27 | 1.00 | 0.00 | 3.13 | 0.00 |
| UB3 | 1.00 | 0.94 | 1.00 | 1.32 | 4.67 | 1.04 | 1.00 | 0.00 | 4.94 | 0.00 |
| UB4 | 1.00 | 0.81 | 1.00 | 0.84 | 5.21 | 0.91 | 1.00 | 0.00 | 5.36 | 0.00 |
| UB | 1.00 | 0.59 | 1.00 | 1.41 | 1.33 | 1.38 | 1.00 | 0.00 | 1.48 | 0.00 |

<sup>a</sup>Selection regime (as in Table 3).
<sup>b</sup>See text for explanation of modified $\mathbf{G}$ matrices.

Substituting 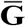 always reduces the change in dimorphism in response to SCS, by an average of 30.4% over our five selective scenarios. Symmetrizing **B** can either increase of decrease the dimorphic response to SCS, but the overall changes in dimorphic response are on average 28.9%, nearly as large as symmetrizing **G**_*m*_ and **G**>_*f*_. Interestingly, the asymmetries due to **B** and **G**_*m*_ and **G**>_*f*_ can have opposite effects. For example, simultaneous SCS on all UB traits leads to smaller changes in dimorphism relative to that under SAS than any other selection regime. The results in Table 4 show that symmetrizing **B** increases dimorphism under SCS by 41%, while symmetrizing **G**>_*m*_ and **G**>_*f*_ decreases dimorphism by 41%. The change in dimorphism is small because the asymmetries work against each other. The predicted response in dimorphism in UB2, however, is large because both types of asymmetry affect dimorphism in the same direction. In addition, the effects of these asymmetries are highly non-linear in combination, as eliminating both kinds of asymmetry complete eliminates the evolution of dimorphism under concordant selection, regardless of how each asymmetry affects dimorphism in isolation. As predicted, symmetrizing the matrices has no effect on the change in dimorphism under SAS. The **B** matrix as a whole is an important constraint to the evolution of dimorphism, as eliminating it always increases the change in dimorphism under SAS.

## Discussion

Most previous analyses of the relationship between sexual dimorphism and genetic variation have made two limiting assumptions (e.g., Poissant, et al. 2010; Griffin, et al. 2013; Matthews, et al. 2019). The first is that sexual dimorphism is shaped by direct, sexually antagonistic selection that favors dimorphism. The second is that genetic constraints on the evolution of dimorphisms can be well-characterized by a single parameter, the intersexual genetic correlation of each trait, *r_MF_*. The latter assumption is as much practical as conceptual: there are relatively few estimates of sex-specific multivariate **G** matrices, compared to the bivariate ones.

We reanalyzed genetic variation in gene expression in *Drosophila melanogaster* (Ayroles, et al. 2009) to include the effects of other types of selection, and to capture the effects of other aspects of inheritance on the evolution of dimorphism. Our major result is that, for this set of gene expression traits, sexually concordant selection (SCS) that selects equally on the phenotypes of each sex is equally capable of causing the evolution of sexual dimorphism in expression as sexually antagonistic selection (SAS) that directly favors dimorphism. In addition, we highlight aspects of genetic variation, other than the values of *r_MF_*, that affect the evolution of dimorphism of gene expression.

Asymmetry in genetic variances between the sexes has long been predicted to cause dimorphism in response to SCS (Fisher 1930; Lande 1980; Leutenegger and Cheverud 1982; Cheverud, et al. 1985;

Lynch and Walsh 1998, Chapter 24; Bonduriansky and Chenoweth 2009; Wyman, et al. 2013), but the role of such differences has rarely been considered as an explanation for observed sexual dimorphisms. An exception is Leutenneger and Cheverud’s (1982) proposal that dimorphism of primate canine teeth and body weight in primates is caused by indirect response to selection on average body size mediated by differences in genetic variances between the sexes.

Our analysis of the two trait case in this paper, as well as a more comprehensive analysis of the *k* trait case (Cheng and Houle accepted) shows that asymmetries in the cross-trait covariances between the male and female genetic variance covariance (**G**) matrices, and asymmetries within the cross-sex covariance (**B**) matrix also play a role in the evolution of dimorphism. The one-trait analyses that feature *r_MF_* omit the role of these aspects of the **G** matrix by assumption. In general, these cross-trait covariances can play a large role in shaping evolutionary trajectories (Blows and Hoffmann 2005; Hansen and Houle 2008; Walsh and Blows 2009). Their omission from existing analyses of gene expression data is particularly unfortunate, as these data give us the opportunity to address effects of indirect selection at an unprecedented scale. Critically, the asymmetries of cross-trait covariances within **G** matrices promote the evolution of dimorphism under concordant selection, and are irrelevant under antagonistic selection. In contrast, the response of dimorphism to antagonistic selection depends only on the average of the male and female **G** matrices, and the symmetrical part of the **B** matrix.

We incorporated these factors into our predictions of dimorphism for *D. melanogaster* gene expression. To generate unbiased estimates of **G**, we analyzed variation in a relatively small number of linear combinations of expression traits, capturing the major axes of variation in female-biased, male-biased and relatively unbiased genes. For 5 of these 8 traits, genetic variation in males and females was highly asymmetrical. This alone will cause the evolution of dimorphism under almost any selection regime. Predictions of response to selection based on the whole **G** matrix showed that sexually concordant and sexually antagonistic selection are approximately equally capable of changing sexual dimorphism. The same result holds for selection that includes non-linear components that restrict the evolution of some traits, while favoring changes in others. If selection in antagonistic and directional selection is equally strong, both will contribute to the level of dimorphism observed in gene expression.

In addition, the indirect responses due to cross-trait correlations were sometimes quite substantial. For example, selection on the highly dimorphic genes is predicted to change the dimorphism in the less-biased genes by a comparable amount to direct selection on the less-biased genes.

We also estimated whether the dimorphism of each gene could be predicted using the relevant gene-specific variances and covariances. As in previous studies, there was strong evidence that between sex-correlations were negatively related to dimorphism, but other aspects of the **G** matrix also explained variation in dimorphism. As predicted under antagonistic selection, the average within-sex genetic variance of the focal trait and average within-sex correlations of the focal trait with all other traits were positively related to dimorphism. Conversely, the difference between male and female trait variances was also positively related to dimorphism, which is expected as a result of indirect responses to sexually concordant selection.

The asymmetries in both of our matrices, **G** and **G\*** are upward biased due to estimation error. The confidence limits for our predicted responses encompass these error, so we are confident that we have detected significant potential for concordant selection to influence dimorphism.

Summarizing these empirical results, analysis of aggregate measures of genetic variation suggests that a substantial amount of variation in dimorphism can be created by directional selection, regardless of whether that selection is antagonistic or concordant. In addition, the dimorphism of individual genes shows traces of both antagonistic and concordant selection.

Lande’s (1980) original explication of the quantitative genetics of dimorphism clearly incorporated the likelihood that concordant selection would affect dimorphism, just as antagonistic selection would. Lande chose to emphasize the effects of antagonistic selection for dimorphism based on the assumption that evolution of sex-averaged means would be relatively unconstrained and rapidly achieve their optima, while evolution of optimal differences between the sexes would tend to be constrained, and take a long time to evolve to their optima. This point is made explicit in the model of Connallon and Clark (2014) who show that the evolution of some dimorphism is almost inevitable whenever selection perturbs any initially monomorphic population. In this case, even if changes in trait optima are random, sex-specific traits will tend to be under antagonistic selection more frequently than they are under concordant selection.

A key motivation for this work is that direct studies of natural selection suggest that, contrary to Lande’s (1980) expectation, there is substantial ongoing concordant selection on traits expressed in both sexes. The available evidence suggests that sexually antagonistic selection is rarer than concordant selection, and more importantly is relatively weak when it is observed (Cox and Calsbeek 2009; Morrissey 2016). The available data on which to base this conclusion is admittedly rather weak (Kingsolver, et al. 2001; Hereford, et al. 2004; Morrissey, et al. 2010; Morrissey and Hadfield 2012). Studies of natural selection tend to be based on rather small sample sizes, use less than ideal measures of fitness, and cannot always account for environmentally induced covariances between traits and fitness (Rausher 1992; Stinchcombe, et al. 2002; Winn 2004). Despite these caveats about the quality of the selection data, if concordant selection is in fact either common or strong, it is clear that we must take seriously the possibility that some sexual dimorphism is just the byproduct of selection for other trait changes.

Lande’s (1980) argument that concordant selection should be rare is based on the assumption that selection regimes change infrequently, allowing the population to approach sex-averaged optima. An alternative world view is that changes in selection are frequent, rather than rare, and populations are rarely at sex-averaged optima. This scenario is consistent with the pattern of sex-specific selection identified by Morrissey (2016), as well as the frequent observation of very strong linear selection gradients in many populations (Kingsolver, et al. 2001; Hereford, et al. 2004).

In conclusion, several aspects of our results suggest the possibility that sexual dimorphism of gene expression in *D. melanogaster* may reflect the indirect effects of concordant selection on the dimorphic traits, or the indirect effects of selection on entirely different traits. Our estimate of the **G** matrix suggest many unexpected correlations and asymmetries that will together generate dimorphism under any selective regime. Strong selection on one aspect of gene expression will frequently generate widespread perturbations in the expression of genes that are not directly selected. The upshot of these factors is that dimorphism in the expression of any particular gene cannot be assumed to be adaptive. We do not doubt that many aspects of transcription do reflect persistent selection for sexual dimorphism. Deciding which aspects of dimorphism are so selected requires more detailed analyses than have so far been applied.

## Acknowledgments

We thank Mark Kirkpatrick, Michael Morrissey, Jacqueline Sztepanacz, Christophe Pélabon, Thomas F. Hansen and members of the Evolvability Project at the Senter for Grunnforskning, Oslo for their discussions of this work. This work was supported by the United States National Science Foundation (www.nsf.gov) Division of Environmental Biology grant 1556774 to D. Houle.

---

**Table S1.**
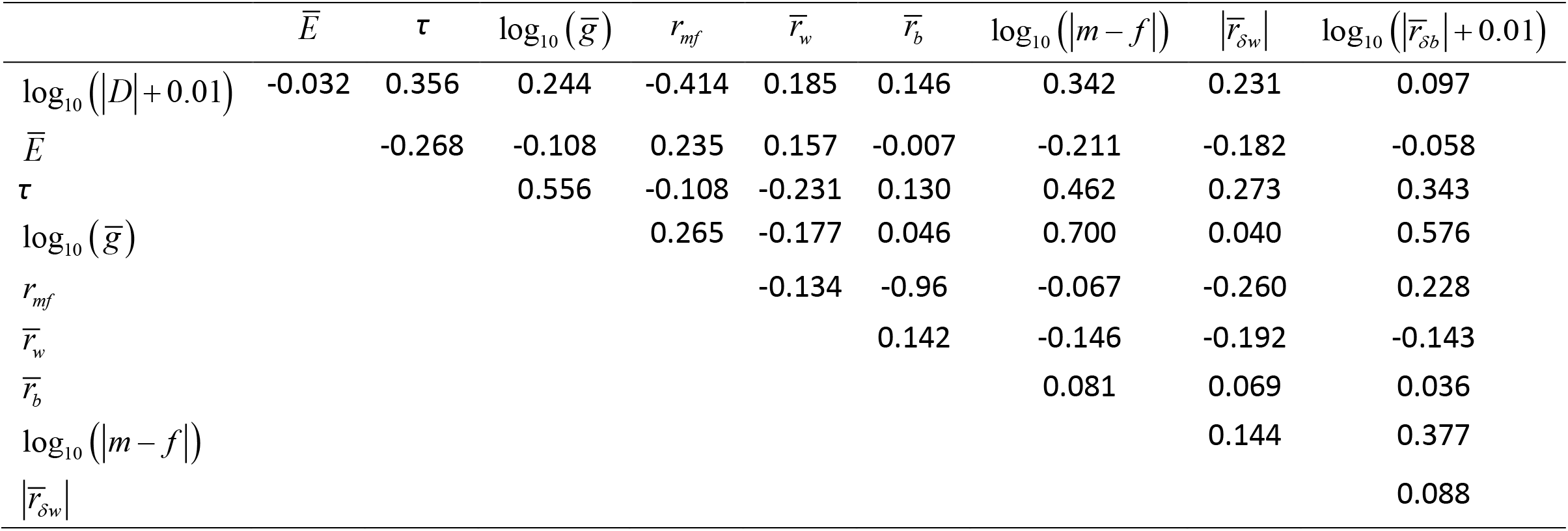
Pearson correlations of dimorphism with predictor variables.

**Table S3.** Genetic variances and standard errors of variance estimates for principal component scores within data subsets defined by degree of sex bias in gene expression for the 16 trait data set.

| Subset | PC | Code | Male |  | Female |  | *d* |
| --- | --- | --- | --- | --- | --- | --- | --- |
|  |  |  | Variance | S.E. | Variance | S.E. |  |
| Male Biased | 1 | MB1 | 53.66 | 14.83 | 6.00 | 4.26 | 0.60 |
| Male Biased | 2 | MB2 | 16.54 | 4.30 | 2.82 | 1.96 | 0.71 |
| Female biased | 1 | FB1 | 5.34 | 2.42 | 88.45 | 22.98 | 0.46 |
| Female biased | 2 | FB2 | 0.29 | 0.12 | 30.62 | 7.31 | 0.19 |
| Unbiased | 1 | UB1 | 21.18 | 4.93 | 102.11 | 23.60 | 0.75 |
| Unbiased | 2 | UB2 | 52.99 | 12.25 | 32.63 | 7.97 | 0.97 |
| Unbiased | 3 | UB3 | 42.02 | 9.93 | 37.72 | 9.13 | 1.00 |
| Unbiased | 4 | UB4 | 33.80 | 8.24 | 39.22 | 9.62 | 1.00 |

**Table S4.** Responses of dimorphism in relatively unbiased gene expression traits to selection on male- and female-biased expression.

| Selection Regime<sup>a</sup> | UB $\ \Delta\ $ <sup>b</sup> |
| --- | --- |
| MB1 | 13.9 (4.3-27.4) |
| MB2 | 4.4 (1.8-8.8) |
| FB1 | 38.3 (21.2-59.1) |
| FB2 | 9.0 (4.5-15.8) |
| MB | 10.7 (3.3-21.7) |
| FB | 25.2 (13.5-40.0) |
| Bias | 10.9 (3.4-21.8) |

<sup>a</sup>MB1, MB2, FB1, FB2 directional selection the indicated trait; MB, FB directional selection on biased traits in one sex; Bias=directional selection on all sex-biased traits simultaneously.
<sup>b</sup>Total change in dimorphism for UB traits. Estimates are mean magnitude of change in dimorphism, (2.5% - 97.5% quantiles).

**Table S5.** Predicted conditional responses (medians and 2.5%-97.5% quantiles) of dimorphism of UB traits to antagonistic (A) or concordant (C) selection when other UB traits are held constant.

| Sel.<sup>a</sup> | Antagonistic<sup>b</sup> | | Concordant<sup>b</sup> | | UB $\ \Delta\ $ <sup>c</sup> | | |
| --- | --- | --- | --- | --- | --- | --- | --- |
|  | *c* | *r* | *c* | *r* | A | C | ratio |
| UB1-c | 8.7 (1.7-17.2) | 15.2 (2.6-32.7) | 36.2 (7.4-71.0) | 38.7 (8.1-75.8) | 6.2 (1.2-12.2) | 8.8 (0.4-20.1) | 0.69 (0.39-3.95) |
| UB2-c | 2.7 (0.7-5.2) | 4.4 (1.4-9.9) | 35.0 (12.2-64.2) | 35.2 (12.2-64.4) | 1.9 (0.5-3.7) | 2.3 (0.1-6.5) | 0.80 (0.20-16.53) |
| UB3-c | 1.6 (0.3-4.1) | 3.6 (0.8-10.2) | 57.6 (27.6-92.8) | 57.8 (27.9-92.9) | 1.2 (0.2-2.9) | 2.1 (0.1-6.9) | 0.51 (0.12-11.77) |
| UB4-c | 1.9 (0.6-3.9) | 3.2 (1.0-7.7) | 51.7 (20.7-86.1) | 51.9 (20.9-86.3) | 1.4 (0.4-2.8) | 1.6 (0.1-5.0) | 0.85 (0.21-17.53) |

<sup>a</sup>Selection regime: the named trait is directionally selected, while the other UB traits are held constant.
<sup>b</sup>Directionally selected male traits have positive gradients, while directionally selected female traits negative selection gradients; *c*=conditional evolvability, the response in the direction of selection; *r*=responsibility, the total response to selection.
<sup>c</sup>Total change in dimorphism for UB traits under A or C selection; ratio= $\|\Delta_A\|/\|\Delta_C\|$ .

